# Dual Roles of Interferon-γ Receptor in Somatostatin Interneurons at Baseline and During

**DOI:** 10.64898/2026.10.06.757042

**Authors:** Sara Rezaei, Toshifumi Tomoda, Michael Marcotte, Thomas D. Prevot, Mounira Banasr, Yashika Bansal, Ali Ezzeddine, Aaron Guan, Erica Vieira, Etienne Sibille

## Abstract

**Background:** Neuropsychiatric disorders such as major depressive disorder show converging evidence of gamma-aminobutyric acid (GABA) dysfunction, including consistent reductions in somatostatin (SST) interneuron markers. SST interneurons are selectively vulnerable to inflammation, yet the underlying mechanisms remain unclear. Interferon-gamma (IFN-γ), a pro-inflammatory cytokine implicated in mood and anxiety disorders, signals through the IFN-γ receptor (IFNGR), which is expressed in neurons. We tested whether IFNGR signaling in SST interneurons regulates anxiety-like and cognitive behavior and prevents inflammation-induced molecular deficits.

**Methods:** We generated mice with conditional deletion of IFNGR from SST cells and assessed anxiety-like behavior using elevated plus maze and working memory using Y-maze at baseline and after lipopolysaccharide (LPS) induced inflammation. Expression of SST-related, neurotrophic, synaptic, and glial markers was quantified.

**Results:** At baseline and after LPS-induced inflammation, IFNGR deletion in SST cells increased anxiety-like behavior without affecting working memory. In control mice, LPS reduced *Sst* (*p=*0.0006), neuropeptide Y (*Npy*) (*p=*0.0022), brain-derived neurotrophic factor (*Bdnf*) (*p*<0.0001), and synapsin 1 (*Syn1*) (p=0.0150); these reductions were not significant in mice lacking IFNGR in SST cells. Knockouts also showed lower baseline *Bdnf, Syn1,* Gephyrin *(Gphn),* and allograft inflammatory factor 1 (*Aif1*, encoding Iba1) than controls.

**Conclusions:** IFNGR signaling in SST cells plays a dual role: supporting normal behavior and molecular homeostasis while mediating LPS-induced molecular deficits. These findings link immune signaling to the GABAergic dysfunction implicated in major depressive disorder and suggest IFNGR signaling in SST cells may regulate cortical microcircuit integrity at baseline and during inflammatory challenge.

## Introduction

Dysfunction of the brain’s principal inhibitory neurotransmitter, gamma-aminobutyric acid (GABA), is a well-established feature of major depressive disorder (MDD). The GABAergic system comprises molecularly and functionally distinct interneuron classes, most prominently somatostatin (SST), parvalbumin (PV) and vasoactive intestinal peptide (VIP) interneurons, along with co-expressed markers such as neuropeptide Y (NPY) and cortistatin (CORT) (1–4). These interneurons form microcircuits that maintain excitation–inhibition balance and support cortical information processing (5). Clinical and postmortem studies consistently report reduced GABA levels (6–9) and decreased expression of SST-related interneuron markers in MDD (10–12). We previously showed that SST interneurons are the GABAergic subtype most vulnerable to peripheral inflammation induced by lipopolysaccharide (LPS) (13), with reduced brain-derived neurotrophic factor (BDNF) contributing to this phenotype (14), although other mechanisms remain uncharacterized. Cytokines are increasingly recognized as mediators of communication between the immune system and cortical GABAergic networks (15,16), exerting neuromodulatory effects on both homeostatic and pathological regulation of neuronal function (17).

Interferon-γ (IFN-γ) is the only type II interferon, produced mainly by natural killer cells and T lymphocytes (18) and, within the central nervous system, by microglia (19). Serum IFN-γ levels are associated with the pathophysiology of MDD (20–22). IFN-γ signals through its heterodimeric receptor (IFNGR), composed of the ligand-binding IFNGR1 subunit and the accessory IFNGR2 subunit (23,24), which activates the Janus kinase/signal transducer and activator of transcription (JAK–STAT) pathway. Activated STAT1 translocates to the nucleus and induces interferon-stimulated genes, including interferon regulatory factor-1 (IRF1), guanylate-binding proteins (GBPs) and suppressor of cytokine signaling-1 (SOCS1) (25). Microglia and astrocytes also express IFNGR and respond to IFN-γ through inflammatory transcriptional programs, suggesting that IFNGR signaling may influence neuroinflammatory processes through neuron–glia interactions (26).

IFNGR is expressed on neuronal soma, dendrites and synapses, supporting a direct role for IFN-γ signaling in neuronal function (27,28). Mice deficient in IFN-γ show increased neurogenesis and improved cognition (29) as well as increased anxiety-like behavior (30,31). IFN-γ also modulates neuronal excitability and inhibitory neurotransmission: it enhances GABAergic neurotransmission *in vitro* (32,33), and conditional deletion of IFNGR from all inhibitory neurons prevents IFN-γ from increasing inhibitory currents (34). During systemic inflammation, IFNGR knockout mice show reduced lethality and cytokine induction following Bacillus Calmette-Guérin and LPS (35,36), and improved memory under chronic stress (37). Together, these findings highlight the importance of distinguishing the physiological function of IFNGR signaling from its pathological effects.

Despite these findings, no studies have directly examined the role of IFNGR signaling within specific subpopulation of GABA interneurons. This gap is particularly important given the convergence of GABAergic dysfunction, SST-specific deficits, and role of inflammation in MDD. In this study, we used a conditional knockout model to investigate how IFNGR in SST interneurons contributes to molecular pathways associated with cortical microcircuit integrity under basal conditions and following LPS-induced inflammation by examining interneuron, synaptic, neurotrophic, and glial markers. In this report, we first hypothesized that IFNGR is expressed on major GABAergic interneuron subtypes in the prefrontal cortex (PFC). Second, based on evidence that disruption of IFN-γ signaling is associated with anxiety-like behavior and cognitive deficits and SST-cell vulnerability, we used mice with a selective deletion of *Ifngr1* in SST cells to test the hypothesis that IFNGR in SST cells contributes to behavioral and molecular homeostasis. Finally, we predicted that selective deletion of IFNGR from SST cells would attenuate the inflammation-induced reductions in SST-related, neurotrophic, synaptic, and glial-associated molecular markers following LPS challenge.

## Methods and Materials

### Animals

Eight-week-old male and female *C57BL/6J* mice (Jackson Laboratories, Bar Harbor, ME) were group housed in individually ventilated cages on a 12 h light/dark cycle with food and water ad libitum. Mice were habituated to the facility for 2 weeks, followed by 3 days of handling (38) to reduce stress towards the experimenter. All procedures followed Canadian Council on Animal Care guidelines and were approved by the Centre for Addiction and Mental Health Animal Care Committee. Mice carrying a floxed *Ifngr1* allele (C57BL/6N-*Ifngr1*^tm1.1Rds/J^, Jackson Laboratories) were crossed with heterozygous *Sst*^tm2.1(cre)Zjh/J^ mice (Jackson Laboratories, stock #013044) to generate *Sst*^cre/+;^ *Ifngr1*^fl/fl^ mice (n=42, 10-14 weeks old, 50% female). *Sst*^cre/+^ littermates served as controls (n=44, 10-14 weeks old, 50% female).

### LPS Injections

Ultra-pure lipopolysaccharide from *Escherichia coli* O111:B4 (InvivoGen, San Diego, CA) was reconstituted in endotoxin-free water, aliquoted in sterile phosphate-buffered saline (PBS) and stored at-20 °C. Mice were weighed and received a single intraperitoneal injection of 2 mg/kg LPS at 3:00 PM, a dose previously shown to be safe and to induce behavioral and molecular changes (39). Based on RNA-sequencing evidence that differentially expressed genes in mouse brain peak one day after LPS (40), tissue was collected at 18 h, a time point that is past the LPS-induced sickness behavior (0-6 hr) (41). Preparation details are provided in the Supplement.

### Quantitative real-time PCR

RNA and protein were extracted from PFC using the AllPrep RNA/Protein kit (catalog no. 80404; Qiagen, Hilden, Germany). cDNA was synthesized from 200 ng total RNA using the SuperScript VILO cDNA Synthesis kit (catalog no. 11754050; Thermo Fisher Scientific, Waltham, MA) in a 20 µL reaction. Amplification used SsoAdvanced Universal SYBR Green Supermix (catalog no. 1725275; Bio-Rad, Hercules, CA) on a Mastercycler real-time PCR system (Eppendorf, Hamburg, Germany), run in triplicate. Primer sets (IDT, Coralville, IA) for all transcripts and for the three internal controls (*Actb*, *Ppia* and *Gapdh*) are listed in Supplemental Table S1. Relative expression was calculated from the geometric mean of the three reference genes and expressed as a percentage of the control group mean; the full calculation is given in the Supplement.

### Enzyme-linked immunosorbent assay

IFNGR1 (catalog no. DY1026) and IFN-γ (catalog no. DY485-05) were measured using DuoSet ELISA kits (R&D Systems, Minneapolis, MN) according to the manufacturer’s instructions. The full assay protocol is provided in the Supplement.

### RNAscope

Fluorescent in situ hybridization was performed to localize mRNA within the PFC using RNAscope Multiplex Fluorescent technology (Advanced Cell Diagnostics, catalog no. 320850). Tissue was taken from C57BL/6 and *Sst*^cre/+;^ *Ifngr1*^fl/fl^ mice given intraperitoneal 2 mg/kg LPS or PBS (2-3 animals per group, 50% female, 9-12 weeks old). Probes targeted mouse *Sst* (404631-C3), *Pvalb (421931-C3)*, *Vip* (415961-C3), *Slc17a7* (416631-C1) and *Ifngr1* (441491-C2); nuclei were counterstained with DAPI. Sections were imaged on an Olympus IX83 microscope with a 60× oil-immersion objective and analyzed in QuPath, quantifying puncta in 50-69 neurons per group from 2-3 mice. Tissue processing and hybridization steps are detailed in the Supplement.

### Elevated-Plus Maze and Y-maze

Anxiety-like behavior was assessed in an elevated plus maze with two open and two closed arms. Mice explored freely for 10 min and video was analyzed with ANY-maze software (Stoelting Co., Wood Dale, IL) to quantify entries into, time spent in, and distance moved in the open arms, and total distance moved in the arena. Apparatus dimensions and lighting are detailed in the Supplement. Spatial working memory was assessed by spontaneous alternation in a three-arm Y-maze. Alternation rate was expressed as a percentage (alternations/trials × 100). The apparatus and full procedure are detailed in the Supplement.

### Statistics

Statistical analyses were performed using GraphPad Prism 10. A three-way ANOVA was initially performed to assess the effects of sex, genotype and LPS, as well as their interactions. As no significant main effect of sex or significant interaction involving sex was detected, data from males and females were pooled. The effects of genotype and LPS treatment were subsequently analyzed using a two-way ANOVA. A power analysis based on the previously observed effect of LPS (Cohen’s f=0.54, α=0.05, power=0.80) indicated that a total sample size of 32 animals, 8 per group, would detect a main effect of LPS in a two-way ANOVA design. When significant main effects or interactions were detected, appropriate post hoc multiple-comparison tests were performed. Outliers were identified using the ROUT method (Q = 1%) (42) in GraphPad Prism and excluded prior to analysis. For RNAscope experiments, differences in puncta per cell between groups were assessed using either unpaired two-tailed Student’s t-tests or Mann-Whitney U tests, as appropriate. Statistical significance was set at *p*<0.05.

## Results

### *Ifngr1* is expressed in SST, PV and VIP interneurons

To determine whether *Ifngr1* is expressed on GABAergic interneurons, we performed multiplex RNAscope in situ hybridization for *Ifngr1* mRNA (red) with the interneuron markers *Sst*, *Vip* or *Pvalb* (yellow) and Dapi counterstain (gray) (Figure 1A-F). Discrete *Ifngr1* puncta localized primarily to the soma of labeled neurons across layers 2-5, demonstrating that *Ifngr1* is expressed across major GABAergic interneuron subtypes in the PFC, including SST, PV and VIP neurons. LPS did not alter *Ifngr1* puncta number in SST (*p=*0.746; Figure 1D) or VIP neurons (*p=*0.199; Figure 1E), whereas PV neurons showed increased *Ifngr1* expression (*p=*0.0365; Figure 1F). Given this baseline expression, the vulnerability of SST interneurons to LPS and their implication in MDD, subsequent experiments focused on IFNGR in SST interneurons.

**Figure 1.**
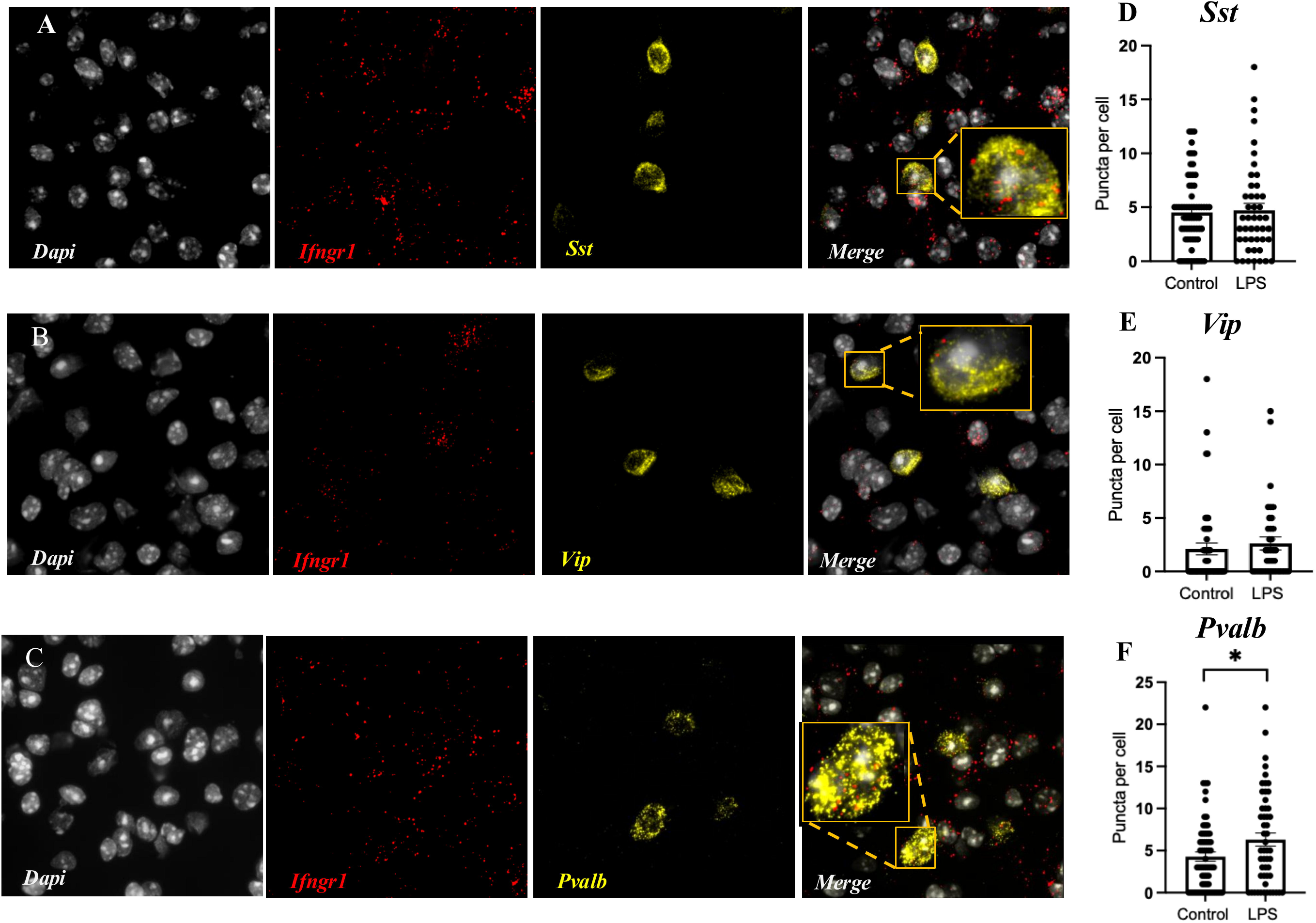
*Ifngr1* expression in cortical SST, VIP, and PV interneurons. (A–C) RNAscope images showing *Ifngr1* mRNA puncta (red) co-labeled with *Sst*, *Vip*, or *Pvalb* (yellow), and nuclei counterstained with *Dapi* (gray) in cortical layers 2–5. Insets show higher-magnification views of representative cells. **(D-F)** Quantification of *Ifngr1* puncta per cell showed no difference between control and LPS groups in SST or VIP interneurons, but a significant increase in PV interneurons following LPS. Data were collected from 2–3 mice per group (control and LPS), with 50–69 neurons analyzed per group. Scale bar, 20 μm in all panels. Each dot represents an individual neuron, bars represent mean ± SEM. \**p* < 0.05.

### *Ifngr1* deletion in SST cells increases baseline anxiety-like behavior without affecting working memory

To investigate the role of IFNGR signaling in SST cells, *Ifngr1* was conditionally deleted in SST cells. Throughout the paper, *Sst*^cre/+^; *Ifngr1*^fl/fl^ mice are referred to as knockouts and *Sst*^cre/+^ mice as controls. RNAscope confirmed the absence of *Ifngr1* in SST-positive cells in knockouts (Figure 2A).

**Figure 2.**
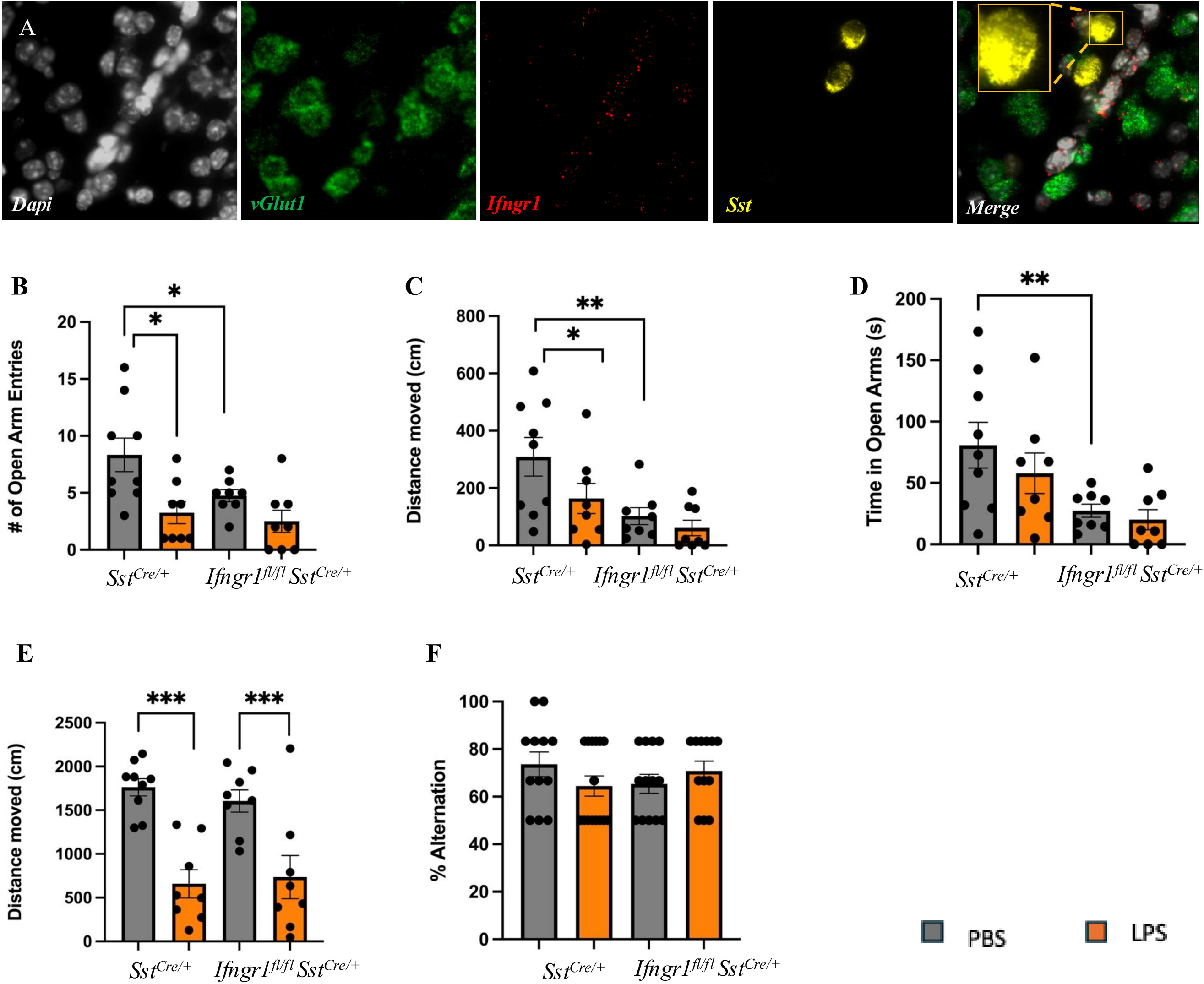
Knocking out *Ifngr1* in SST interneurons increased baseline anxiety-like behavior without affecting working memory. (**A**) RNAscope images showing *Ifngr1* mRNA (red), *vGlut1* (Green), co-labeled with *Sst* (yellow), and nuclei counterstained with *Dapi* (gray). Expression of *Ifngr1* was not detected in SST cells in mice lacking IFNGR in SST neurons. Scale bar, 20 μm in all panels. Following intraperitoneal injection of PBS or LPS (2 mg/kg, 18 h), anxiety-like behavior was assessed in the elevated plus maze (EPM): **(B)** number of open arm entries, **(C)** distance moved in open arms, **(D)** time spent in open arms, and **(E)** total locomotor activity, measured as distance moved in the arena. **(F)** Spatial working memory assessed by Y-maze. For elevated plus maze *n* = 33; 8-9 mice/group (B–E) and *n* = 52; 12-15 mice/group for Y-maze (F). Data are shown as individual animals with mean ± SEM. \**p* < 0.05, \*\**p* < 0.01, \*\*\**p* < 0.001.

In the elevated plus maze (EPM), open-arm entries showed a main effect of LPS (*F₁,₂₉=* 11.50, *p=*0.002), a trend toward a main effect of genotype (*F₁,₂₉=* 4.015, *p=*0.0545), and no interaction. At baseline, knockouts made fewer open-arm entries than controls (*p=*0.0242); LPS reduced entries in controls (*p=*0.0021) but not in knockouts (*p=*0.1576; Figure 2B).

Distance moved in the open arms showed a main effect of genotype (*F₁,₂₉=* 10.01, *p=*0.0036), a trend toward a main effect of LPS (*F₁,₂₉=* 3.652, *p=*0.0659), and no interaction. At baseline, knockouts traveled a shorter distance than controls (*p=*0.005); LPS reduced distance in controls (*p=*0.0413) but not in knockouts (*p=*0.5603; Figure 2C).

Time spent in the open arms showed a main effect of genotype (*F₁,₂₉=*10.89, *p=*0.0026), with no main effect of LPS and no interaction. At baseline, knockouts spent less time in the open arms than controls (*p=*0.0097); LPS did not alter this in controls or knockouts (Figure 2D).

Total distance moved in the arena, a measure of locomotor activity, showed a main effect of LPS (*F*₁,₂₉=35.78, *p*<0.0001), with no main effect of genotype and no interaction. LPS reduced locomotor activity in both controls (*p*<0.0001) and knockouts (*p=*0.001; Figure 2E).

Spatial working memory, assessed by spontaneous alternation in the Y-maze, showed no main effect of genotype (F₁,₄₈=0.043, p=0.8368), no main effect of LPS (F₁,₄₈=0.176, p=0.6770), and no interaction (F₁,₄₈=2.715, p=0.1060; Figure 2F).

Overall, *Ifngr1* deletion in SST cells produced anxiety-like features at baseline that resembled those seen after LPS, without further potentiation by LPS, while working memory was unaffected. We next examined molecular markers of GABAergic interneurons, glial and synaptic markers.

### *Ifngr1* deletion attenuates LPS-induced reductions in SST interneuron markers

We next measured mRNA levels of *Sst*, *Npy*, *Cort*, *Pvalb* and *Vip*. *Sst* showed a main effect of LPS (*F*₁,₃₇=12.28, *p=*0.0012), with no main effect of genotype and no interaction. Baseline *Sst* tended to be lower in knockouts than controls (*p=*0.09). LPS reduced *Sst* in controls (*p=*0.0006) but not in knockouts (*p=*0.1622; Figure 3A).

**Figure 3.**
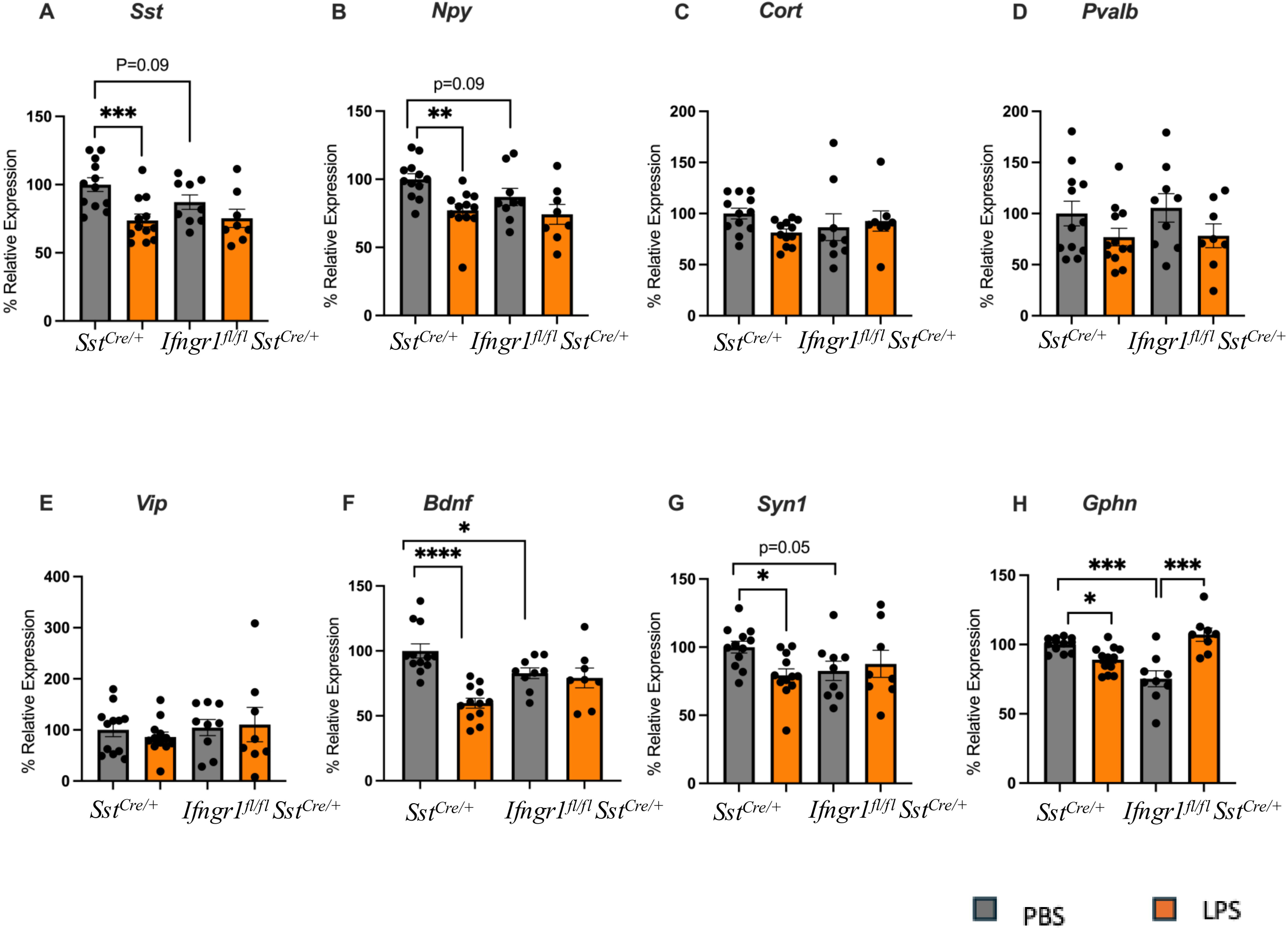
Expression of GABAergic interneuron markers, *Bdnf, Syn1* and *Gphn* in control and mice lacking *Ifngr1* in SST cells. (A–H) Relative mRNA expression of GABAergic interneuron markers (*Sst, Npy, Cort, Pvalb, Vip*), neurotrophic (*Bdnf*), and synaptic markers (*Syn1, Gphn*) in control mice (*Sst*^Cre/+^) and mice lacking *Ifngr1 (Sst*^cre/+;^ *Ifngr1*^fl/fl^) in SST cells following PBS or LPS treatment (2 mg/kg, 18 h);. Data are shown as individual mice (n=41; 8-12 mice/group) with mean ± SEM. \**p* < 0.05, \*\**p* < 0.01, \*\*\**p* < 0.001, \*\*\*\**p* < 0.0001.

*Npy* showed a main effect of LPS (*F*₁,₃₇=10.91, *p=*0.0021), with no main effect of genotype and no interaction. Baseline *Npy* also tended to be lower in knockouts (*p=*0.09). LPS reduced *Npy* in controls (*p=*0.0022) but not in knockouts (*p=* 0.1306; Figure 3B).

*Cort* and *Vip* showed no effect of genotype or LPS (Figure 3C, E). *Pvalb* showed a main effect of LPS (*F*₁,₃₇=4.604, *p=*0.0385), with no significant post hoc effect in controls (*p=*0.1357) or knockouts (*p=*0.1367; Figure 3D).

### *Ifngr1* deletion reduces baseline *Bdnf*, *Syn1* and *Gphn* and alters their LPS responses

*Bdnf* showed a main effect of LPS (*F*₁,₃₇ =17.24, *p=*0.0002) and a genotype × LPS interaction (*F*₁,₃₇=12.01, *p=*0.0014), with no main effect of genotype. At baseline, knockouts had lower *Bdnf* than controls (*p=*0.0247); LPS reduced *Bdnf* in controls (*p* <0.0001) but not in knockouts (*p=*0.656; Figure 3F).

*Syn1* showed a genotype × LPS interaction (*F*₁,₃₇=4.190, *p=*0.0478), with no main effects. Baseline *Syn1* tended to be lower in knockouts (*p=*0.055). LPS reduced *Syn1* in controls (*p=*0.0150) but not in knockouts (*p=*0.5994; Figure 3G).

*Gphn* showed a main effect of LPS (*F*₁,₃₇=8.199, *p=*0.0069) and a genotype × LPS interaction (*F*₁,₃₇=34.23, *p*<0.0001), with no main effect of genotype. At baseline, knockouts had lower *Gphn* than controls (*p*<0.0001). LPS reduced *Gphn* in controls (*p=*0.0258) but increased it in knockouts (*p*<0.0001; Figure 3H).

### *Ifngr1* deletion decreases baseline *Aif1*; LPS increases *Gfap*

We next measured the astrocytic and microglial markers *Gfap* and *Aif1* (encoding Iba1). *Gfap* showed a main effect of LPS (*F*₁,₃₇=34.38, *p*<0.0001), with no main effect of genotype and no interaction. Baseline *Gfap* did not differ between genotypes (*p=*0.654); LPS increased *Gfap* in both controls (*p=*0.0005) and knockouts (*p*<0.0001; Figure 4A).

**Figure 4.**
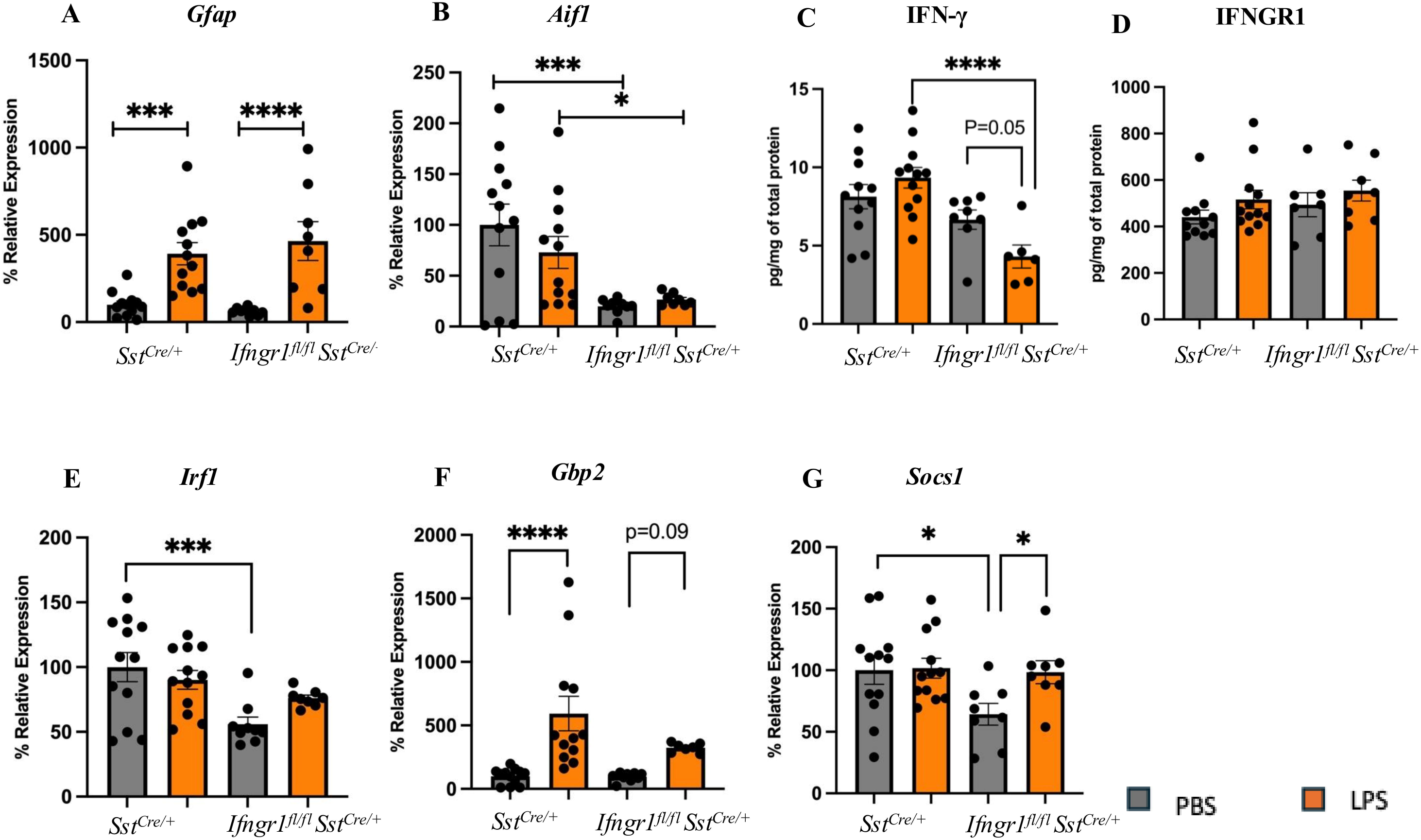
Expression of *Gfap, Aif1*, IFNGR1, IFN-γ, *Irf1, Gbp2,* and *Socs1* in control mice and mice lacking *Ifngr1* in SST cells (A-B) Relative mRNA expression of *Gfap* and *Aif1* in control mice (*Sst*^Cre/+^) and mice lacking *Ifngr1 (Sst*^cre/+;^ *Ifngr1*^fl/fl^) in SST cells following PBS or LPS treatment (2 mg/kg, 18 h). **(C-D)** Protein levels of IFN-γ and IFNGR1 in control mice (*Sst*^Cre/+^) and mice lacking *Ifngr1 (Sst*^cre/+;^ *Ifngr1*^fl/fl^) in SST cells following PBS or LPS treatment (2 mg/kg, 18 h). (**E-G**) Relative mRNA expression of *Irf1*, *Gbp2,* and *Socs1*. Data are shown as individual mice (n=37-41; 6-12 mice/group) with mean ± SEM. \**p* < 0.05, \*\**p* < 0.01, \*\*\**p* < 0.001, \*\*\*\**p* < 0.0001.

*Aif1* showed a main effect of genotype (*F*₁,₃₇ =29.72, *p=*0.0002), with no main effect of LPS and no interaction. Knockouts had lower *Aif1* than controls at baseline (*p=*0.0007) and after LPS (*p=*0.0453); LPS itself did not alter *Aif1* in controls (*p=*0.184) or knockouts (*p=*0.782; Figure 4B).

### *Ifngr1* deletion reduces *Irf1* and *Socs1*; LPS increases *Gbp2*

We next measured IFN-γ, IFNGR1, and the downstream signaling factors *Irf1*, *Gbp2* and *Socs1*. IFN-γ showed a main effect of genotype (*F*₁,₃₃=18.53, *p=*0.0001) and a genotype × LPS interaction (*F*₁,₃₃=5.56, *p=*0.0244), with no main effect of LPS. Baseline IFN-γ did not differ between genotypes (*p=*0.1622). LPS did not alter IFN-γ in controls (*p=*0.1993) and tended to reduce it in knockouts (*p=*0.0575), such that after LPS IFN-γ was lower in knockouts than controls (*p*<0.0001; Figure 4C). IFNGR1 showed no main effect of LPS or genotype and no interaction, consistent with deletion restricted to a sparse interneuron population that is not detectable in bulk cortical tissue (Figure 4D).

*Irf1* showed a main effect of genotype (*F*₁,₃₇ =11.88, *p=*0.0014); at baseline, knockouts had lower *Irf1* than controls (*p=*0.0006; Figure 4E).

*Gbp2* showed a main effect of LPS (*F*₁,₃₆=17.77, *p=*0.0002), with no main effect of genotype and no interaction. LPS increased *Gbp2* in controls (*p*<0.0001) and tended to increase it in knockouts (*p=*0.0941; Figure 4F).

*Socs1* showed trends toward main effects of genotype (*F*₁,₃₆=3.782, *p=* 0.0597) and LPS (*F*₁,₃₆ = 3.185, *p=*0.0827). At baseline, knockouts had lower *Socs1* than controls (*p=*0.0165); LPS increased *Socs1* in knockouts (*p=*0.0348; Figure 4G).

Overall, *Ifngr1* deletion in SST cells lowered baseline expression of neurotrophic, synaptic, microglial, and interferon-responsive markers and changed how these markers responded to inflammation. LPS reduced *Sst, Npy, Bdnf* and *Syn1* in controls, and these reductions were absent in knockouts, while *Gphn* decreased in controls but increased in knockouts. In contrast, LPS-induced *Gfap* and *Gbp2* expression was retained in both genotypes, indicating that the inflammatory challenge remained biologically effective.

## Discussion

To investigate a potential mechanism underlying the selective vulnerability of SST neurons to inflammation, we examined the role of IFNGR in SST neurons and found that cell-type specific deletion of *Ifngr1* disrupts baseline behavioral and molecular homeostasis, while providing some degree of protection against LPS-induced molecular deficits. Together, these results support a dual, context-dependent role for IFNGR signaling in SST interneurons.

### Constitutive role of IFNGR in SST interneurons

*Ifngr1* is constitutively expressed in cortical inhibitory interneurons, consistent with transcriptomic datasets reporting *Ifngr* expression in mouse hippocampal interneurons (32,43). To determine the functional relevance of this expression, we assessed behavior in mice with conditional deletion of *Ifngr1* in SST cells. Because IFN-γ must bind IFNGR1 to initiate receptor complex formation, deletion of IFNGR1 effectively abolishes IFN-γ signaling. At baseline, knockouts displayed increased anxiety-like behavior without changes in locomotor activity, indicating a specific effect on affective rather than motor function, whereas working memory was unchanged. Similarly other studies found that global knock out of the *Ifngr1* at baseline also increased emotionality such as increased anxiety-like behavior (30,31,44) and reduced social behavior (34). Another study found that at baseline, mice lacking IFN-γ showed impaired spatial memory measured by Y-maze (37). Since we do not find any changes in memory function when *Ifngr1* is knocked out in SST cells, it suggests that constitutive IFNGR signaling in SST neurons is more relevant to affective rather than cognitive domains.

Baseline behavioral alterations were accompanied by reduced *Irf1* and *Socs1* expression, suggesting that constitutive IFNGR signaling in SST cells regulates selected interferon-responsive pathways in the PFC. Molecular changes followed the same pattern: trends toward reduced *Sst* and *Npy*, consistent with impaired maintenance of SST-related molecular identity, and a significant reduction in baseline *Bdnf*. Notably, BDNF was not reduced in global IFN-γ knockout studies (31,37), suggesting that SST-specific deletion alters cortical microcircuit activity and the local neurotrophic environment, producing secondary changes in BDNF expression from surrounding neuronal and glial populations. Synaptic markers followed suit: the postsynaptic GABA_A_ receptor scaffold *Gphn* was significantly reduced at baseline, and the presynaptic *Syn1* showed a comparable trend (45,46). Given that IFN-γ enhances GABAergic inhibitory transmission through inhibitory interneurons (34), the coordinated reductions across SST-related, synaptic and neurotrophic markers suggest that loss of IFNGR signaling in SST interneurons weakens SST/GABAergic activity and inhibitory synaptic maintenance within the local cortical microcircuit.

Consistent with the notion that loss of IFNGR signaling alters the cortical microcircuit, SST-specific deletion of *Ifngr1* also reduced baseline *Aif1* expression. Microglia are sensitive to both neuronal activity and neurotrophic signaling, and changes in inhibitory circuit function as well as reduced *Bdnf* expression may influence neuron–microglia communication (47–51). In contrast, baseline *Gfap* expression was not altered by SST-specific deletion of *Ifngr1*, suggesting that physiological IFNGR signaling in SST interneurons does not substantially influence astrocytic homeostasis under basal conditions. Thus, the coordinated reductions in SST-associated markers, synaptic genes, *Bdnf*, and *Aif1* suggest that physiological IFNGR signaling contributes to maintaining not only SST interneuron integrity but also broader microcircuit–microglia homeostasis within the cortex.

These baseline effects differ from global IFN-γ or IFNGR knockout models, where developmental alterations and broader immune deficits complicate interpretation. By deleting *Ifngr1* selectively from SST cells while preserving IFN-γ signaling elsewhere, our approach isolates a cell-type-specific contribution of IFNGR signaling to maintaining adult affective behavior and molecular integrity, rather than secondary consequences of global immune disruption or altered development.

### IFNGR signaling contributes to LPS-induced molecular vulnerability

Having established that IFNGR signaling in SST cells contributes to baseline behavioral and molecular homeostasis, we next examined whether the same pathway mediates the molecular vulnerability of SST cells during systemic inflammation. In control mice, LPS-induced inflammation significantly reduced the expression of *Sst, Npy, Bdnf,* and the synaptic markers *Syn1* and *Gphn*. In contrast, these LPS-induced reductions were attenuated in mice that lacked the *Ifngr1*, indicating that deletion of *Ifngr1* from SST cells reduces inflammation-induced molecular deficits.

These results align with evidence that IFN-γ signaling mediates the biological effects of LPS. IFNGR-deficient mice show reduced systemic responsiveness to endotoxin, including resistance to LPS-induced anorexia, hypermetabolism and BCG-potentiated lethality (36,52). IFN-γ blockade or STAT1 deficiency prevents LPS-induced inflammatory activation in microglia and human monocytes (53,54), and in slice cultures LPS and IFN-γ act synergistically: neither alone is sufficient to cause neuronal death, whereas combined exposure triggers iNOS-dependent nitric oxide release and cell death (55).

Analysis of IFN-γ-related markers further showed that deletion of *Ifngr1* in SST cells modified the local response to LPS. IFN-γ protein showed a significant genotype x LPS interaction, with markedly lower levels in LPS-treated knockout compared to LPS-treated controls; LPS did not alter IFN-γ in controls but produced a trend toward reduced IFN-γ in knockouts. This suggests that IFNGR signaling in SST cells influences IFN-γ regulation during systemic inflammation, although whole-PFC measurements do not identify the cellular source. That total IFNGR1 protein was unaltered at 18 h may reflect the transient, time-dependent nature of cytokine signaling and activation through pre-existing receptor complexes (56–58). Nevertheless, LPS increased *Gbp2*, a well-established interferon-stimulated effector gene, across genotypes, indicating a broader interferon-responsive program that remained largely intact following *Ifngr1* deletion. The induction of *Socs1* by LPS in knockouts further suggests that inflammatory negative-feedback mechanisms remained responsive.

Notably, LPS did not increase *Aif1* expression in either genotype, whereas *Gfap* was robustly induced in both, indicating that the inflammatory challenge remained biologically effective and that *Ifngr1* deletion did not globally suppress glial responses.

Together, these findings position IFNGR signaling in SST cells as a mediator of inflammation-induced molecular vulnerability rather than a driver of the inflammatory response itself.

### A dual role for interferon gamma receptor signaling

A central question is how IFNGR deletion produces impairments at baseline while reducing molecular deficits during LPS-induced inflammation. This is consistent with IFN-γ signaling operating within an inverted U-shaped relationship with neuronal and behavioral homeostasis, as proposed for other brain cytokines (59,60): both insufficient and excessive signaling are detrimental, whereas tightly regulated physiological signaling supports optimal function.

Interleukin-1 (IL-1) is the canonical example: physiological signaling is required for memory and modest elevations enhance learning, whereas either excessive activation or blockade impairs cognition (60–65). IFN-γ is likewise present at low, tightly regulated levels in the central nervous system (66), contributing to normal neuronal and behavioral regulation (67), and transgenic models of sustained low-level IFN-γ expression show enhanced neurogenesis, neuroprotection and improved cognition, attributed to immune–glial support of neurotrophic pathways including BDNF (68,69). Consistent with this, deletion of *Ifngr1* from SST interneurons reduced baseline *Bdnf*.

On the left of the curve (Figure 5), insufficient signaling is detrimental. Global IFNGR knockout mice show increased anxiety-like and depressive-like behavior, cognitive impairment and reduced neurogenesis (30,31,37,44), and deletion of STAT1 from GABAergic neurons induces social deficits and disrupts IFN-γ–mediated regulation of tonic and phasic inhibitory currents (34). Knockouts here map onto this side under basal conditions, displaying behavioral and molecular deficits alongside reduced basal *Irf1* and *Socs1*, indicating disrupted constitutive interferon-responsive signaling rather than induction of a neuroprotective state.

**Figure 5.**
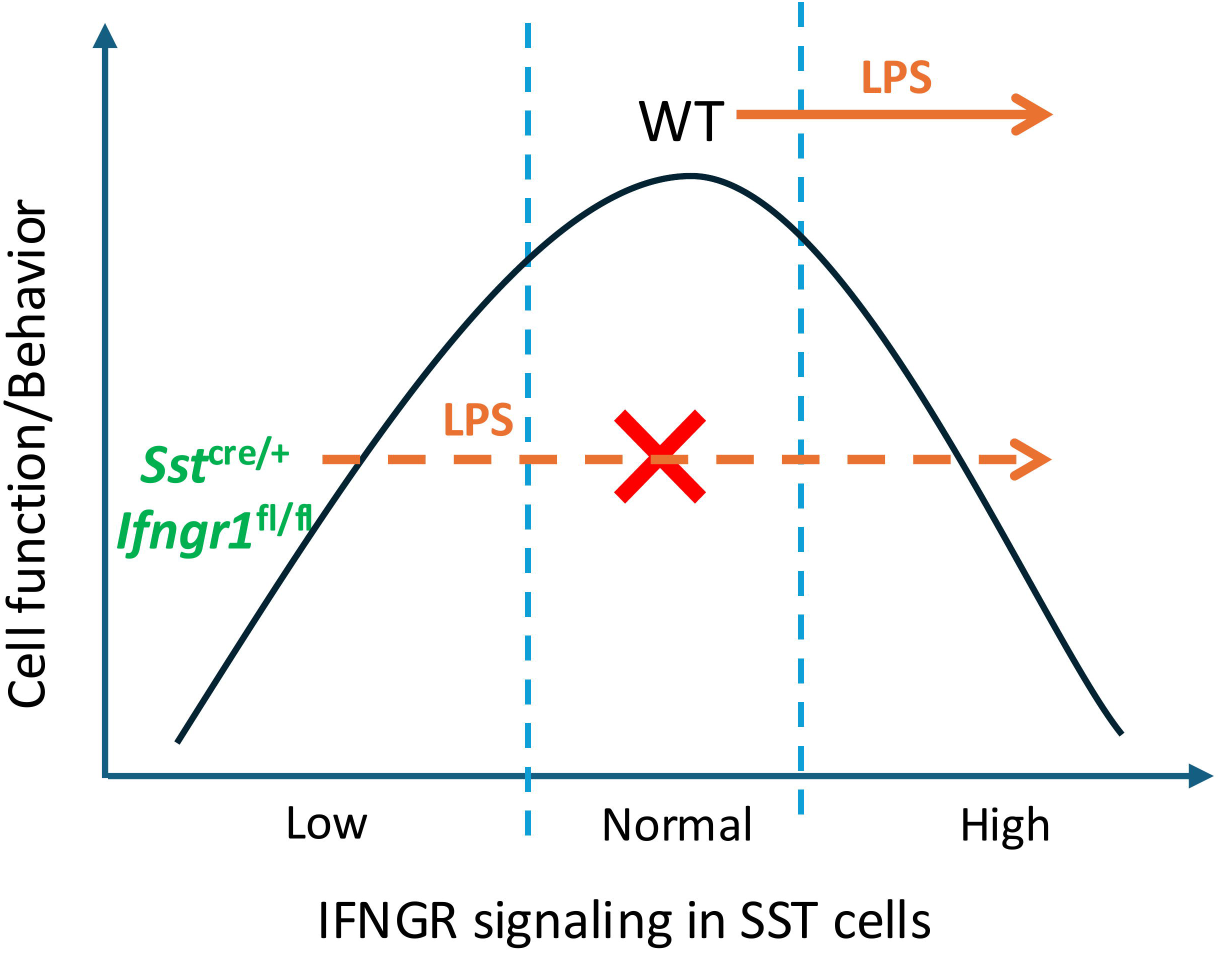
Inverted U-shaped relationship between IFNGR signaling in SST cells and cell function/behavior. Schematic illustrating the proposed dual role of IFNGR signaling in SST interneurons. Under physiological conditions, IFNGR signaling support molecular and behavioral homeostasis in wild-type (WT) mice. Insufficient IFNGR signaling, as in SST-specific *Ifngr1* knockout mice at baseline is associated with reduced neurotrophic, synaptic markers and increased anxiety-like behavior. During LPS-induced inflammation, IFN-γ signaling is shifted toward excessive activation in WT mice, contributing to inflammation-induced molecular vulnerability. Deletion of *Ifngr1* prevents this pathological rightward shift, preserving SST, neurotrophic and synaptic related molecular markers.

On the right of the curve, excessive or prolonged IFNGR activation, as during systemic LPS exposure, becomes maladaptive and contributes to altered neuronal excitability and neurotoxicity (26,27,70–72). Duration and magnitude appear critical: physiological signaling induces brief, tightly regulated STAT1 activation, whereas pathological exposure produces prolonged signaling and sustained transcriptional change (73,74). Consistent with this, chronic IFN-γ reduced GABAergic inhibitory tone (75), high-dose acute IFN-γ increased hippocampal excitability (76), and prolonged IFN-γ disrupted network oscillations (77).

Together, these findings support a dual, context-dependent role for IFNGR signaling in SST cells. Under basal conditions, physiological signaling maintains neurotrophic and synaptic support and regulates affective behavior. During systemic inflammation, LPS may shift signaling toward the maladaptive range, suppressing SST markers, *Bdnf* and synaptic gene expression. Deletion of *Ifngr1* prevents this shift and preserves those markers during inflammation, but does not restore an optimal physiological state: it mitigates selected LPS-induced deficits while producing impairments at baseline.

### Limitations

Although our findings are consistent with the proposed U-shaped model (Figure 5), this non-linear relationship was not directly tested. Future studies incorporating a broader range of affective, cognitive and social behavioral tests would help further define the functional consequences of SST-specific *Ifngr1* deletion. Finally, our hypothesis-driven approach on IFNGR and a panel of molecular markers was necessarily limited in scope. Unbiased approaches such as single-cell RNA sequencing, could identify additional immune receptors and pathways regulating specific GABAergic interneuron subtypes.

## Acknowledgments

SR is supported by University of Toronto Labatt Family Doctoral award, a funding that ES received from the Labatt Family Network for research on the Biology of Depression. Research was also supported by CAMH Discovery Fund (EV). MB is supported by the CAMH Discovery Seed Fund and the Canadian Institutes of Health Research (PJT-165852). We would like to thank CAMH animal facility personnel. During the preparation of this work, the authors used OpenAI’s ChatGPT to assist with language editing of author-written text. The authors reviewed and edited the output and take full responsibility for the content of the publication.

