## Supplement 1 for "Dual Roles of Interferon-γ Receptor in Somatostatin Interneurons at Baseline and During"

**Dual Roles for Interferon-γ Receptor Signaling in Somatostatin Interneurons at Baseline and During Inflammation**

**Supplemental Methods**

**Quantitative real-time PCR**

RNA concentration was measured by NanoDrop spectrophotometry. Each qPCR reaction was run in triplicate in 96-well white-shell/clear-well plates. The three internal control genes were beta-actin (*Actb*), cyclophilin A (*Ppia*) and glyceraldehyde-3-phosphate dehydrogenase (*Gapdh*). Cycle threshold (Ct) values were used to calculate delta Ct (dCt) against each reference gene, and the geometric mean of the three dCt values was taken. Relative expression level was calculated as 2^-dCt × 1,000, then converted to a percentage by dividing each value by the mean relative expression level of the control group.

**Enzyme-linked immunosorbent assay**

Total protein extracted from PFC was quantified by bicinchoninic acid assay (Pierce BCA Protein Assay Kit, catalog no. 23225; Thermo Fisher Scientific, Waltham, MA) according to the manufacturer's instructions. A 96-well microplate was coated with 100 µL per well of diluted capture antibody in PBS and incubated for 18 h. Plates were washed manually three times (300 µL per well) with wash buffer (1× PBS with 0.1% Triton X-100), blocked with 300 µL of reagent diluent (1% BSA in PBS) for 2 h, then washed three times. Cytokine standards and samples diluted 1:2 were added and incubated for 18 h at 4 °C. After washing, 100 µL of detection antibody was added and incubated for 2 h. Following a further wash, 100 µL of streptavidin-HRP was added and the plate incubated for 30 min at room temperature. After a final wash, 100 µL of TMB substrate solution was added for 20 min, followed by 50 µL of stop solution. Optical density was read at 450 nm on a BioTek ELx800 absorbance reader (Agilent Technologies, Santa Clara, CA).

**RNAscope**

Brain tissue was immediately flash-frozen, sectioned at 20 µm on a Leica CM1950 cryostat, mounted on SuperFrost Gold Plus slides (Thermo Fisher Scientific, catalog no. 1255015) and stored at -80 °C until analysis. Sections were fixed in 4% paraformaldehyde and dehydrated through a graded ethanol series (50%, 70% and 100%), then treated with Protease IV to make mRNA accessible for hybridization. Hybridization was carried out using the HybEZ Hybridization System. Following hybridization, sections underwent signal amplification with Amp1-4 reagents and nuclei were counterstained with DAPI. Images were analyzed in QuPath, an open-source platform for digital pathology image analysis, which enabled cell detection and subcellular localization for quantification of target gene expression within SST, PV and VIP cells.

**Y-maze**

The Y-Maze apparatus was constructed from black polyvinyl chloride (PVC) and consisted of three arms arranged at 120° angles, forming a Y-shaped configuration. Each arm was equipped with a sliding door at its entrance to control access during trials. Mice were first habituated to the maze by being allowed to explore freely for 10 minutes per day over two consecutive days. On the pretesting third day, animals underwent training to alternate between arms. Each trial began with the mouse placed in the start box for a 30-second acclimation period before the door was opened, allowing it to choose freely between the left or right arm. Once an arm was selected, the mouse was confined within that arm for 30 seconds before being gently returned to the start box for a 30-second inter-trial interval (ITI). This procedure was repeated for 7 trials. If an animal failed to alternate for three consecutive trials, a forced alternation trial was introduced to avoid reinforcement of side preference. On the testing day, fourth day, the same general procedure used in the training session is implemented 18 h later, except that the ITI is lengthened to 60 and an 8^th^ trial is added with a shorter ITI (10 sec) to test motivation to alternate in an easy paradigm. The mean alternation rate is calculated and is expressed in percentage (number of alternation/number of trials × 100) as an index of cognition. Mice that fail to alternate in the 8^th^ trial were excluded from analysis. Each mouse was tested twice in the Y-Maze: once prior to LPS injections to assess baseline performance and again following the 18 h LPS injection.

**Elevated-Plus Maze**

Animals were individually tested in the elevated plus maze (EPM) to assess anxiety-like behavior. The apparatus comprised four Plexiglas arms arranged in a plus-shaped configuration, consisting of two opposing open arms (67 × 7 cm) and two opposing enclosed arms (67 × 7 × 17 cm). The maze was elevated 95 cm above the floor and placed in a dimly lit testing room (approximately 20 lux). At the start of each trial, mice were positioned in the central square facing an open arm and allowed to explore freely for 10 minutes. Behavioral activity was recorded from an overhead camera mounted on a fixed rod. Video recordings were subsequently analyzed using ANY-maze tracking software (Stoelting Co., Wood Dale, IL, USA) to quantify the time spent and number of entries into the open and closed arms.

**Supplemental Table S1**

*Table S1. Primer sequences used for quantitative real-time PCR*. All primers were synthesized by IDT (Coralville, IA).

| **Gene** | **Primer 1 (5′–3′)** | **Primer 2 (5′–3′)** |
| --- | --- | --- |
| *Sst* | CAACTCGAACCCAGCAAT | GGTCTGGCTAGGACAACAA |
| *Npy* | TTAATGAAGGAAAGCACAGAAA | AGATGAGATGAGGGTGGAAA |
| *Cort* | CCTTCTCCTCGTGCAAGTA | AGGTCTCGTTGGCATCTC |
| *Pvalb* | ATGAGGTGAAGAAGGTGTTCC | AGCGTCTTTGTTTCTTTAGCAG |
| *Vip* | GACATCTTGCAGAATCCCTTA | CTGCTGTAATCGCTGGTG |
| *Bdnf* | CAGTATTAGCGAGTGGGTCA | CCTTTGGATACCGGGACT |
| *Gphn* | GAGACAACCCAGATGACTTACTC | GAAGATCAATGTCCAGCACCT |
| *Gfap* | TGCAGGAGTACCAGGATCTAC | GATCTGGAGGTTGGAGAAAGTC |
| *Aif1* | GAACCCTCTGATGTGGTCTG | AGGAGGACTGGCTGACTT |
| *Actb* | CCTAGCACCATGAAGATCAA | GGAAGGTGGACAGTGAGG |
| *Ppia* | CAAACACAAACGGTTCCCAG | TTCACCTTCCCAAAGACCAC |
| *Gapdh* | AACTCCCACTCTTCCACC | CACCACCCTGTTGCTGTA |
| *Irf1* | GTCACCCATGCCTTCCAC | ACTCAGACTGTTCAAAGAGCTTC |
| *Gbp2* | GCAGAATTCACCTCATACATCTTG | GATGGCACCAACATAGGTCTG |
| *Socs1* | ATTCCACTCCTACCTCTCCAT | CAGAAAAATGAAGCCAGAGACC |
| *Ifngr1* | GCTTCTTTGAAGGGCTGGA | TTGTTTGATCCATTCTTTAAATTG |
